# Conjugation factors controlling F-plasmid antibiotic resistance transmission

**DOI:** 10.1101/271254

**Authors:** Hanna Alalam, Fabrice E. Graf, Martin Palm, Marie Abadikhah, Martin Zackrisson, Matilda Mattsson, Chris Hadjineophytou, Linnéa Persson, Simon Stenberg, Payam Ghiaci, Per Sunnerhagen, Jonas Warringer, Anne Farewell

## Abstract

The rapid horizontal transmission of many antibiotic resistance genes between bacterial host cells on conjugative plasmids is a major cause of the accelerating antibiotic resistance crisis. Preventing understanding and targeting conjugation, there currently are no experimental platforms for fast and cost-efficient screening of genetic effects on antibiotic resistance transmission by conjugation. We introduce a novel experimental framework to screen for conjugation based horizontal transmission of antibiotic resistance between >60.000 pairs of cell populations in parallel. Plasmid-carrying donor strains are constructed in high throughput. We then mix the resistance plasmid carrying donors with recipients in a design where only transconjugants can reproduce, measure growth in dense intervals and extract transmission times as the growth lag. As proof-of-principle, we exhaustively explored chromosomal genes controlling F plasmid donation within *E. coli* populations, by screening the Keio deletion collection at high replication. We recover all six known chromosomal gene mutants affecting conjugation and identify >50 novel factors, all of which diminish antibiotic resistance transmission. We verify 10 of the novel genes’ effects in a liquid mating assay. The new framework holds great potential for exhaustive disclosing of candidate targets for helper drugs that delay resistance development in patients and societies and improves the longevity of current and future antibiotics.

## INTRODUCTION

Antibiotic resistance (ABR), particularly in Gram-negative bacteria, is an accelerating crisis. In 2014, most areas of the world reported greater than 50% of *Escherichia coli* infections being resistant to 3rd generation cephalosporins, widespread resistance to fluoroquinolones and accelerating resistance to 3rd generation carbapenems (WHO, 2014). Only few new antibiotics against Gram-negatives are in clinical trials and the pipeline is not predicted to be large enough to keep up with the rate of resistance emergence (WHO, 2017). New approaches to this problem are therefore sorely needed. A major problem is that many antibiotic resistance genes can be transmitted horizontally into and between human pathogens (Norman, et al, 2009). Horizontal transmission within pathogenic species, combined with the selective pressure imposed by extensive antibiotic use, subsequently facilitates their extremely rapid spread. The drastic decline in clinical potency of most of our frontline, as well as ‘last-resort’ antibiotics, including cephalosporins, carbapenems, and most recently, colistins, is predominantly due to horizontal transmission of antibiotic defense factors (Canton, et al, 2013, Carottoli, 2013) - and most often via plasmid conjugation. The transferred conjugative elements can be then maintained as plasmids or integrated into the host chromosome (ICEs, integrative conjugative elements).

Plasmids are self-replicating genetic modules capable of dissemination through conjugation and, to a lesser extent, transformation (Frost, et al 2005, Halary, et al, 2010, Lacroix, et al, 2016, Norman, et al 2009). More than 6000 proteobacterial plasmids have been sequenced (NCBI, 2018) and the association of different conjugative plasmid families with various antibiotic resistances have been studied in *Enterobacteriaceae* (Carattoli, et al, 2009). Conjugation involves production of a pilus (encoded by the conjugative element) that attaches to target cells and facilitates the transfer of the conjugative element to the recipient. It has recently been suggested that an effective approach to limit the spread of ABR would be to inhibit conjugation of resistance-carrying plasmids (Banquereo, et al, 2011, Fernandez, et al, 2005, Garcillan-Barcia, et al, 2007, Getino, et al, 2016, Lin, et al, 2011), by chemically blocking conjugation factors in either donors or recipients. However, plasmid encoded conjugation factors are not always well conserved across plasmids, decreasing their value as drug targets. Conjugative elements vary in host range, suggesting that plasmid donation or reception often also depends on chromosomally encoded factors in donors and recipients.

Few of these chromosomal genetic determinants of conjugation are known because of the absence of an approach that is sufficiently fast and cost-efficient for unbiased screening of tens of thousands of bacterial populations. Measuring conjugation efficiency generally relies on slow, meticulous mating assays that are prohibitively expensive and labour intensive to scale-up. Moderate throughput designs were introduced to screen for conjugation effects in recipient cells, but disclosed only a few effects (eg, Perez-Mendoza and de la Cruz, 2009). Here, we develop, implement and validate a massive throughput experimental framework to monitor the conjugation of resistance carrying plasmid donor libraries to a recipient cell population in near real time. We expect the framework to accommodate screening of a wide variety of clinically relevant plasmids, species and environments and to become invaluable in the search for chemical inhibitors of conjugative spread of ABR.

## RESULTS

### A high throughput platform for measuring conjugation of antibiotic resistance plasmids

We designed a platform capable of measuring the conjugative transmission of antibiotic resistance at massive throughput. We robotically construct *E. coli* donor strains and then collect donor and recipient cells from distinct source plates and deposit them as a mixed population on a plate that is double selective for two non-interfering, bacteriostatic antibiotic resistances (Fig 1A). Antibiotic resistance is chromosomally encoded in recipient cells but plasmid encoded in donors. Therefore, only recipient cells that have received a plasmid from a donor, i.e. transconjugants, will divide on the double selective plate. The growth lag of the mixed population will reflect the time to conjugate the plasmid and express its resistance gene. Fixing the recipient genotype, the expression time becomes a constant and conjugation time variation equals lag time variation. To measure lag time, we adopted a recently introduced platform, Scan-o-matic (Zackrisson, et al, 2016), for surveying *S. cerevisiae* colony population size expansion at a massive scale. We deposited 1152 mixed populations on each plate, maintained plates on flatbed scanners in thermostatic cabinets and acquired transmissive light images every 10 minutes. Colonies are identified and background subtracted and pixel intensities are extracted and transformed into cell counts in a fully automated procedure. (Fig 1A). First, we mated *E. coli* donor cells carrying an F plasmid with tetracycline resistance to *E. coli* recipients with chloramphenicol resistance in a conjugation neutral loci (Δ*araB*::cam) in 768 mixed populations (Fig 1B). We obtained an average lag-time of approximately 5.46 h (at 30°C), with only small spatial variation across the plate (Coefficient of variation = 19%). Pure donor and pure recipient cell populations uniformly failed to grow (Fig 1B). An *E. coli* strain with both resistance markers grew with no detectable lag-phase (Fig 1C).

**Figure 1.**
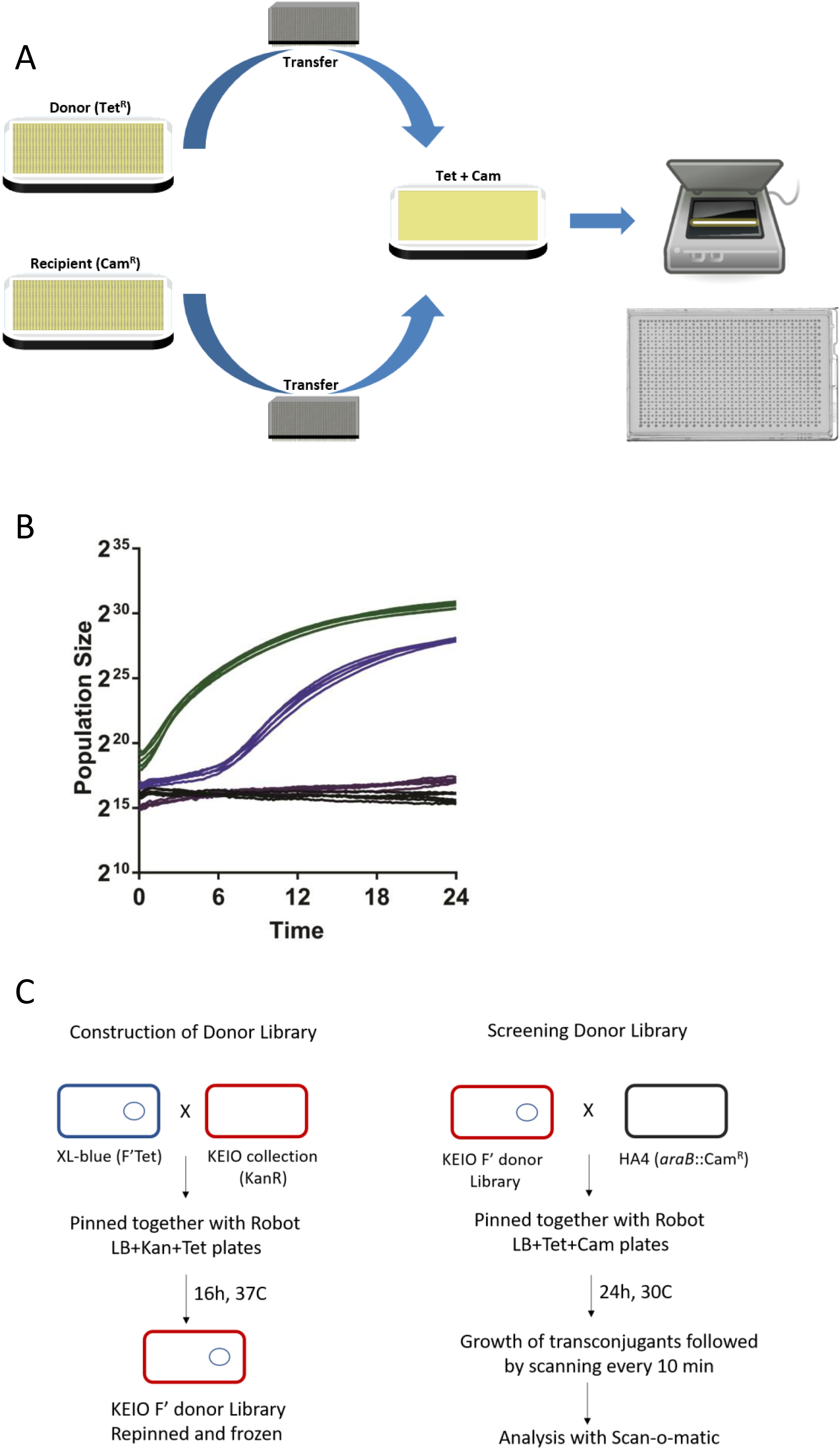
Experimental scheme for screening strains for conjugative efficiency. **A Robotic setup** Donor (F’ Tet^R^) and recipient (chromosomal Cam^R^) strains are grown separately on appropriate pre-culture plates and then pinned robotically in a 1536 format to a selective plate (Tet Cam) that only allows transconjugants to grow. Plates are placed in a flatbed scanner and scanned every 10 minutes for 24 hours at 30°C. Data is then analysed with Scan-o-matic as described in the text. A representative agar plate after incubation is shown. **B Growth of Transconjugants formed on Selective Plates** Blue: Growth of spots pinned with HA4 (chromosomal Cam^R^) and HA14 (F’Tet^R^) together showing growth of the resulting transconjugants which occurs after a lag as compared to Green: HA5 (Cam^R^ Tet^R^) which grows with no detectable lag. Negative controls: violet, HA4 alone and black, HA14 alone are shown. 5 representative graphs are shown for each. **C Construction and Screening of Donor Library** Donor library was constructed by mating XL1-Blue with the Keio collection carrying a kanamycin resistance gene in place of almost 4000 genes is shown on the right. Subsequent screening of the conjugation efficiency is shown on the left. Mating was then performed and analysed as indicated in Figure 1A and the text.

### Comprehensive view of donor functions controlling *F* plasmid conjugation

Next, we introduced the tetracycline resistance carrying F plasmid into the 3908 deletion strains of the *E. coli* KEIO library (Baba, et al, 2006) by mating to a fixed XL1-Blue genotype carrying F’Tet and multiple rounds of double selection (Fig 1C). We subsequently mated the Keio donor library to a fixed recipient genotype (HA4; chloramphenicol resistance) at moderate replication (*n*=8; on two plates), while monitoring the conjugation using the massive throughput platform (Zackrisson, 2016). We deposited 1152 populations on each plate, interleaving 384 genetically identical controls (HA14 x HA4) in every 4^th^ position to control for any systematic spatial effects. We extracted the lag time for each experimental population, normalized it to that of neighbouring controls and expressed the ratio on a log_2_ scale. Positive numbers reflect long lag time and delayed conjugation compared to the control. Overall, donor gene effects on conjugation were symmetrically distributed (*μ*=-0.062, *σ*=0.23) around the WT mean, with extremes being more common than expected from a normal distribution and somewhat more likely to correspond to delayed conjugation (Fig 2A; Fig S1). We selected 60 of the worst affected gene deletions for further validation as the most promising drug targets, as well as 28 weaker hits down to rank 240 to test reproducibility also for more marginal effects. We validated these 88 candidates in a high replication (*n*=48) secondary screen (Fig 2C; Suppl Table 1) along with 6 very low ranked candidates as negative controls. Gene effects on conjugation generally agreed well (r^2^=0.56) between the primary and secondary screen with 81 of the 88 strains chosen from the first screen giving longer lag times than the control mating (Suppl Table 1; Fig 2B). Specifically, we recovered all previously described chromosomal mutants known to affect F plasmid (or F-like plasmids) conjugation (*arcA, crp, hda dnaK, dnaJ, ihfA* (Fig 3A, Beutin and Achtman, 1979, Starcic, et al, 2003, Modrzewska, et al, 2002, Kato and Katayama, 2001, Williams and Schildback, 2007). We also confidently identified 50 novel genes whose deletions were defective in F plasmid donation in both assays. The top 50 encoded gene products covered a wide range of cellular functions, but were disproportionately likely to mediate DNA replication (6 proteins, *p*<10^-5^), chaperone or protein folding functions (6 proteins, *p*<10^-5^) and lipopolysaccharide core biosynthesis (4 proteins, *p*<0.001) (Fisher’s exact test, EcoCyc, Keseler, et al, 2017) (Fig 3 and Table S1). We considered three sources of confounding effects. First, *rimM* and *rnt* were initially scored as conjugation deficient, but were discarded as likely false positives due to their very poor growth on the background LB medium (they do not form detectable single colonies on LB medium in 24 hours). Second, cross-referencing our candidates with chloramphenicol hypersensitive deletions (Liu, et al, 2010): in our secondary screen, we recovered 5 out of 19 mutants classified as strongly chloramphenicol sensitive (*tolC, flgF, rfaG, rfaE* and *acrB*), thus these would need to be independently verified in the absence of the antibiotic (*rfaE* was verified, below). Finally, we discarded one top candidate, *yjjY*, because it is a deletion of a small ORF that overlaps the *arcA* promoter (Compan and Touti, 1994).

**Table 1.**
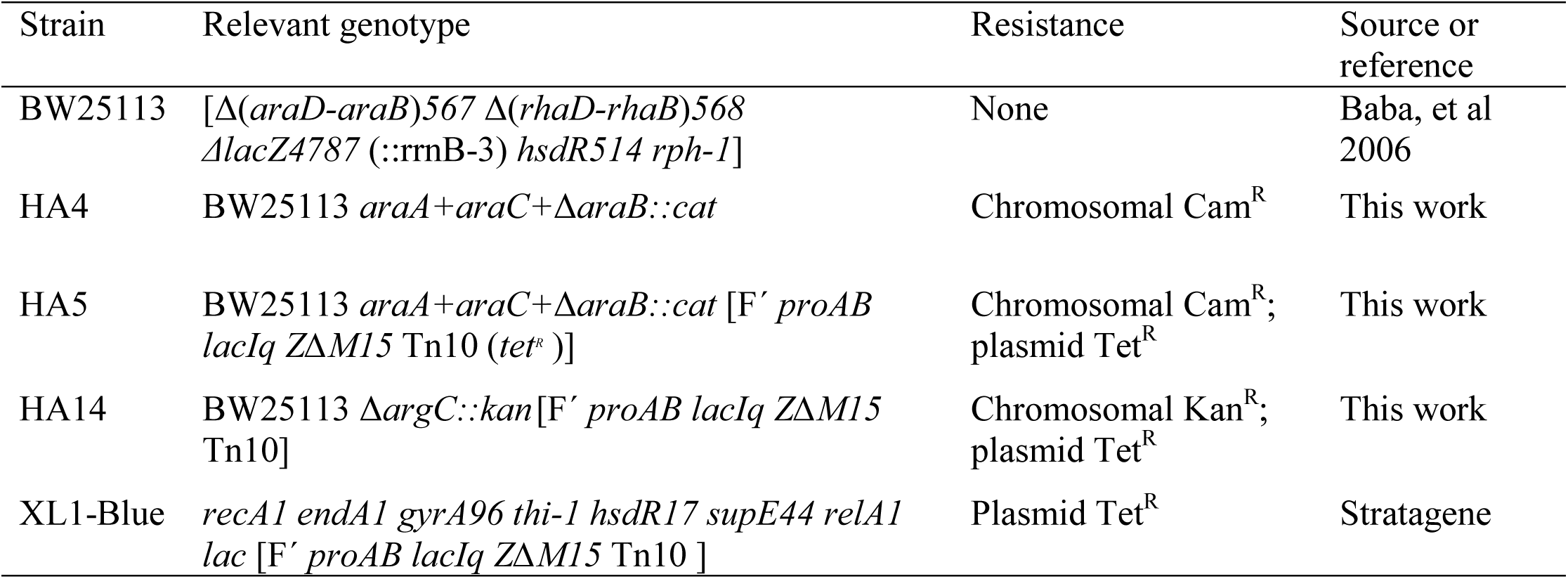
*E. coli* strains used in this work.

**Figure 2.**
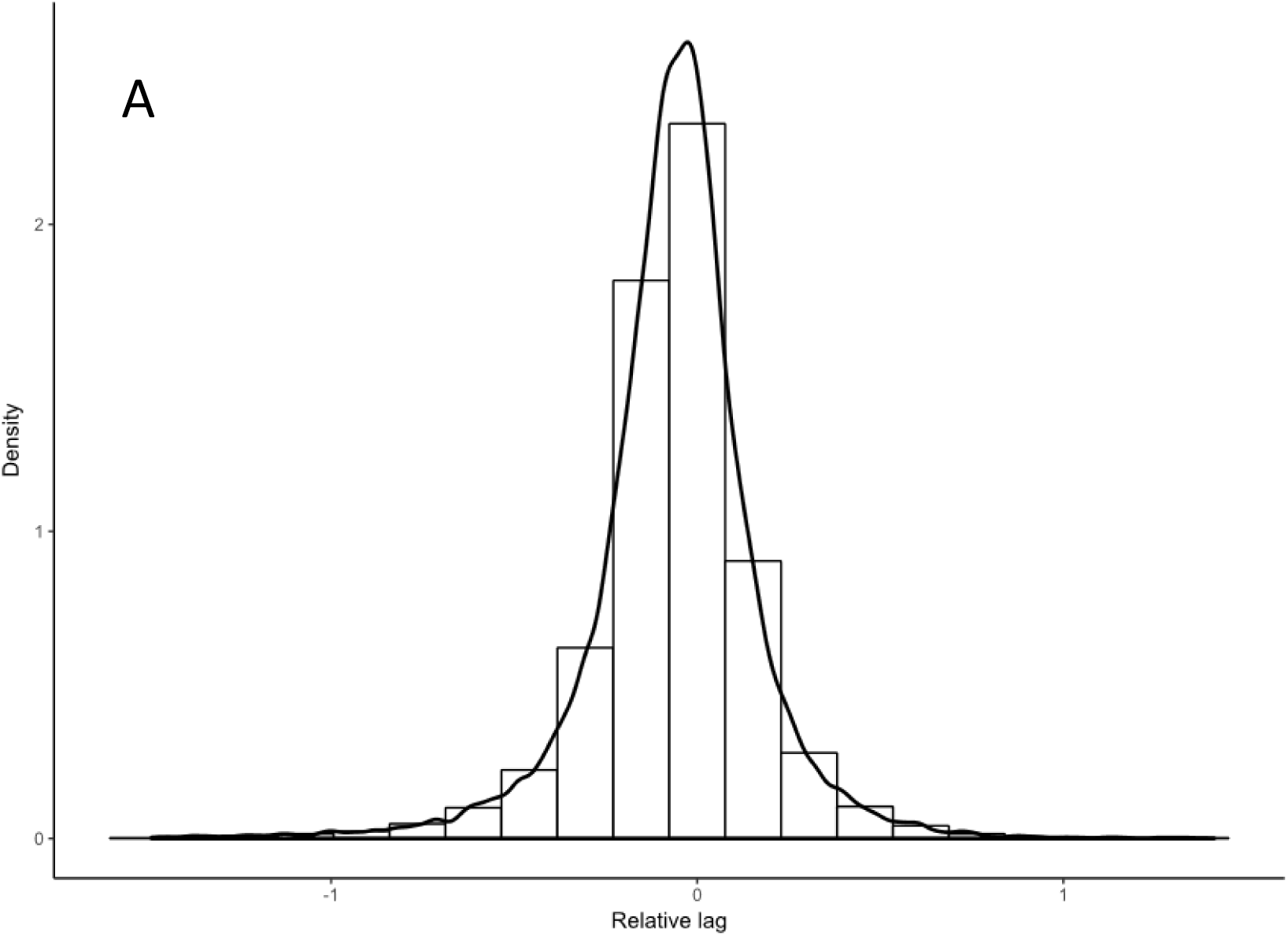
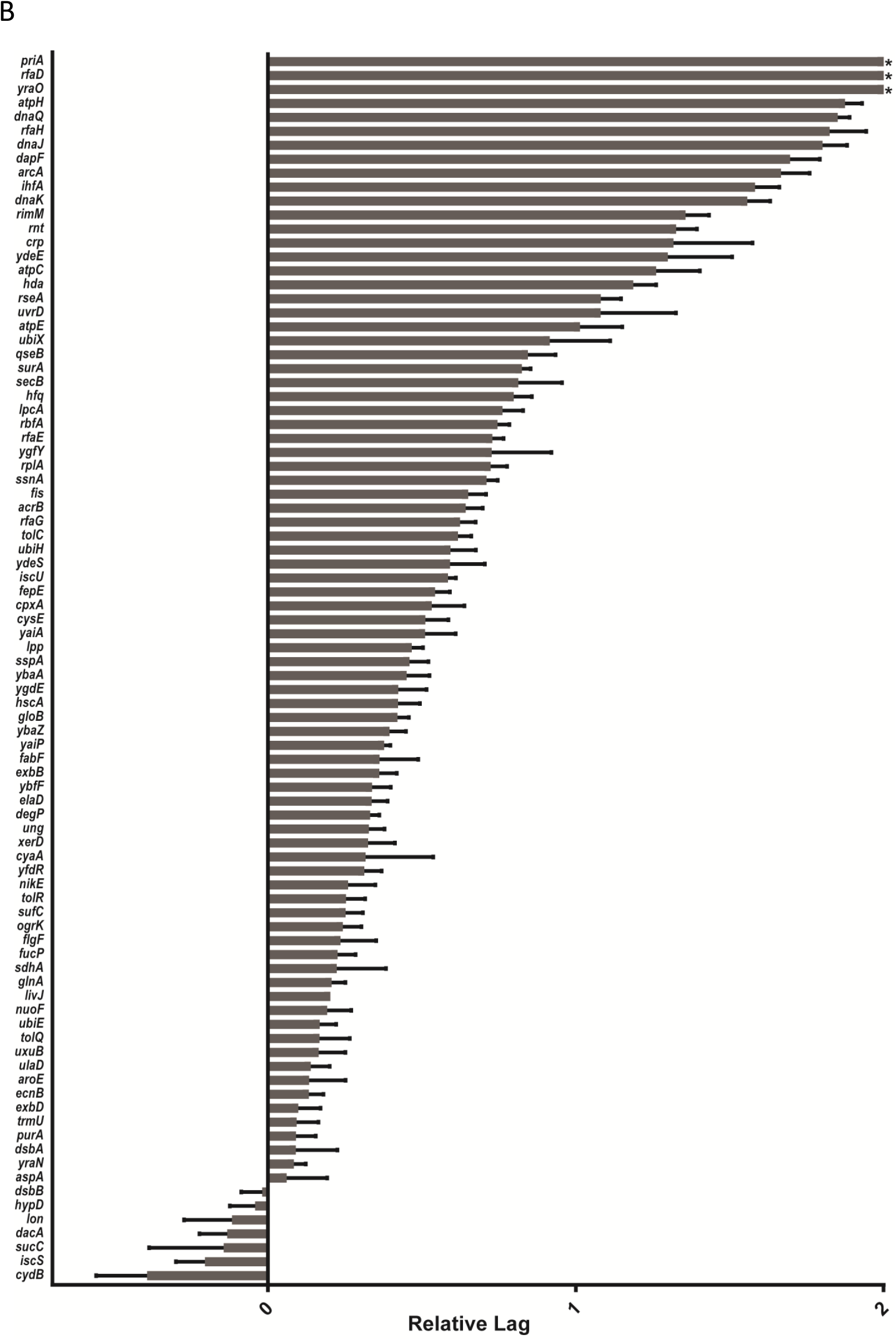
Conjugative Efficiency of the Keio collection. **A Growth lag time** of the entire Keio collection. The growth lag of each curve is expressed relative to the nearby control and the log_2_ value calculated. Shown is the frequency plot for the collection: a positive value indicates that the strain has a longer lag period and a negative number indicates it is shorter than the control mating. **B Growth lag time of the top candidates** 88 strains from the first screen (Figure 2A) were selected and the conjugation efficiency screening repeated with 48 replicates. Strains were chosen as described in the text. Plotted is the mean of the values with the standard error of the mean. Especially in the top candidates, many of the replicates had no detectable conjugation. For comparison we have set values in this dataset with no measurable growth lag to a value of ‘2’ which corresponds to a mating lag time of four times the local control. Three of the strains had fewer replicates (*yraO, ygfY* and *crp* had 8, 8, and 12 replicates respectively). ‘*’ indicates that none of the replicates had measurable lag times.

**Figure 3.**
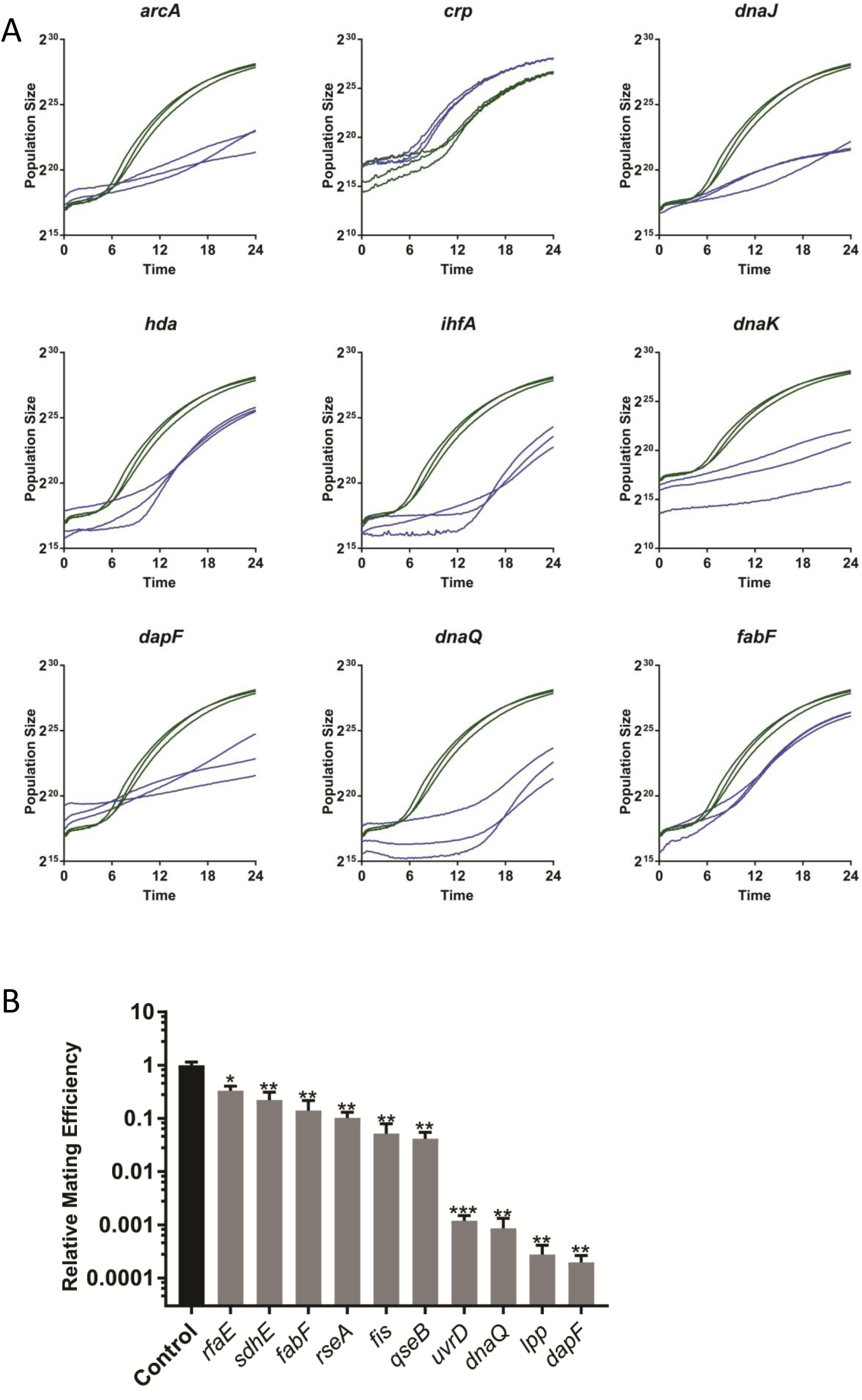
Conjugative efficiency from strains with detectable defects in mating. **A Representative growth curves** from 6 previously known and three newly identified conjugation deficient mutants. The deleted gene in each strain is indicated. Three curves were taken from the plate screening experiments for each mutant as indicated (green) and three nearby control mating results (blue). **B Liquid mating assay results** from 10 newly identified conjugation deficient strains. The gene deleted in each mutant is indicated and the average mating efficiency (number of transconjugants per number of donor cells divided by the corresponding value of a control mating done on the same day) in a 30 minute liquid mating assay with 4-6 biological replicates is shown. Error bars indicate the standard error of the mean. P values were calculated using a one-sided student t-test (* indicates p<0.05, ** indicates p<0.01 and *** indicates p<0.001).

To exclude confounding cross-contamination and strain construction errors, we validated the absence of the expected gene in 36 of the top conjugation deficient deletion by PCR; all were confirmed to be deleted (not shown). We next validated the conjugation defects of 10 mutants in an independent, liquid mating assays. Unlike the plate assays, conjugation in liquid mating assays occurred in the absence of antibiotics. Because the distinction between liquid and solid medium conjugation is important, with liquid matings often detecting additional effects (i.e., mating pair stabilization, Arutyunov and Frost, 2013), we did not expect a direct correlation (e.g., compare mating deficiency in the *lpp* mutant (Fig 3B vs Fig S2). Nevertheless, all 10 mutants showed conjugation deficiencies also in liquid, ranging from 0.02 to 33% of the conjugation efficiency of the wild type (Fig 3B). Four of these (*dapF, lpp, dnaQ, uvrD*) had very strong defects, with near absent conjugation. DapF, encoding a diaminopimelate (DAP) epimerase that leads to *meso*-DAP formation in the peptidoglycan (Antia et al., 1957; Richaud et al., 1987), and Lpp, a membrane lipoprotein that interacts with *meso*-DAP (Dramsi et al., 2008), are particularly interesting drug targets; DAP analogs already exist but have not yet been tested for anti-conjugative properties (Baumann et al.,1988; Lam et al., 1988; Gelb et al., 1990; Gerhart et al., 1990; Gillner et al., 2009). The other two very strong, confirmed candidates are both involved in DNA repair or replication: UvrD is a DNA helicase involved in DNA repair and has been implicated in the replication of some plasmids that use rolling circle replication (Lestini and Michel, 2007) (unlike F which uses θ replication (Bruand and Ehrlich, 2000)) and DnaQ is a subunit of DNA polymerase III (Echols, et al 1973). The role of these two mutants in F conjugation is unknown but may indicate unexplored factors in F replication and conjugative transfer. Among the 10 mutants, we also saw significant effects even in the mutants with the weakest effect on conjugation (*rfaE* and *sdhE*) (*p*<0.01, Fig 3B); this indicates that mutants ranked down to position 74 in our initial screen are promising candidates for conjugation defects and should not be disregarded.

### Discussion

We designed a high throughput platform for measuring conjugation of antibiotic resistance plasmids and demonstrated its utility by identifying all previously known *E. coli* genes that control F plasmid donation, as well as >50 novel conjugation genes not previously linked to antibiotic resistance transmission. The novel conjugation deficient mutants spanned an unexpectedly wide range of functions. Some of these can be rationally explained, e.g., those altering the cell surface including lipopolysaccharide. Lipopolysaccharide mutants have previously been reported to have defects in acting also as conjugation recipients (Perez-Mendoza, et al 2009) and may affect the mating pair interaction. The effect of *dapF* and *lpp* mutants will be explored further to determine if they are deficient in assembly or stabilisation of the F pilus. UvrD and DnaQ likely affect the replication of the F plasmid (directly or indirectly). Other mutants have no obvious connection to conjugation and likely act indirectly, e.g., by controlling expression (transcription, translation), folding (chaperone) and energy supply for conjugation components.

IncF are narrow host-range (limited to *Enterobacteriaceae*) plasmids but are highly diverse within their group and associated with ESBL-producing *E. coli*; an incFII containing CTX-M15 ESBL (Extended Spectrum Beta-Lactamase gene) was likely a contributor to the emergence and establishment of the globally dominant *E. coli* sequence type 131 (ST131) (Novais, et al, 2007, Nicolas-Chanoine et al, 2014)). Nevertheless, IncF plasmids are not the only plasmids of high clinical concern, with Inc A/C, L/M, N, I1 and HI2 plasmids all representing major challenges (Carattoli, 2013). Whether the same or distinct chromosomal factors control transmission of non-IncF plasmids is unknown but critical to any drug development effort. We note that no conceptual challenges prevents identifying the determinants of Inc A/C, L/M, N, I1 and HI2 plasmid transmission, using the introduced platform.

The methodology we have developed is highly adaptable for similar experimental designs targeting conjugation in other bacteria, or cross-species conjugation. Strong coloration in bacteria or background medium or massive secretion of polysaccharides can interfere with correct population size estimations and will to some degree affect conjugation time estimates; we have encountered no other factors that constrain the general applicability of the platform.

## ACKNOWLEDGEMENTS

Funding for this project was provided by grants from the Centre for Antibiotic Resistance Research at the University of Gothenburg to AF and JPIAMR TransPred project (SRC grant number 2016-06503_3) provided to JW. The Keio strain collection was ordered from National BioResource Project (NIG, Japan):E.coli. The students of the course BIO510 at GU from 2017 and 2018 are gratefully acknowledged for their input into this work. Sofia Hultgren and Owens Uwangue are acknowledged for technical support and helpful discussions.

## MATERIALS AND METHODS

### Strains and media

LB medium was routinely used (5 g/l yeast extract, 10 g/l tryptone and 10 g/l NaCl, 15 g/l agar was added if needed). When appropriate, the medium was supplemented with chloramphenicol (30 µg/ml), kanamycin (50 µg/ml), tetracycline (10 µg/ml) hereafter referred to as Cam, Kan and Tet, respectively. Liquid cultures were grown in a rotary shaker at 37°C at 220 rpm. Strains are listed in Table 1. Strain HA4 was constructed by replacing *araB* in MG1655 with a Cam resistance marker using primer FWD araB CAM (ATTGGCCTCGATTTTGGCAGTGATTCTGTGCGAGCTTTGGCGGTGGACGTGTAGGCTGGAGCTGCTTC) and primer RVS araB CAM (AAGTTGGA AGATAGTGTTGTTCGGCGCTCATCGCCCATTGCTGATAGCGATGGGAATTAGCCATGGTCC) to amplify the Cam gene from pKD3, followed by transformation into xxx and P1 transduction into BW25113 as previously described (Datsenko and Wanner, 2000). Strain HA5 was constructed by conjugating XL1-Blue with HA4 on solid LB for approximately 3 hours and then selecting for transconjugants on LB Tet Cam. Strain HA14 was retrieved from the donor library (see below) and streaked on LB Tet Kan.

### Initial testing of the system

Frozen 96-well stock plates of HA4 (Cam^R^ recipient), HA14 (control donor F’Tet^R^) and HA5 (control Cam^R^ Tet^R^ strain) were created by mixing an overnight culture with glycerol to a final concentration of 15% and adding 175 µl to each well. Pre-cultures were prepared by pinning from the 96-well plates to positions as described in Fig 4. Pre-cultures were incubated at 30°C for 16 hours. Subsequently, the control HA5 pre-culture was transferred to a LB Tet Cam plate (pre-warmed to room temperature) using a 1536-short pad (Singer LTD, UK) followed by the recipient pre-culture and then the donor pre-culture to the same LB Tet Cam plate resulting in positions containing matings of HA4x HA14, negative controls and control growth positions (HA5), (see fig 4). The pinned plate was then moved to an Epson Perfection V800 PHOTO scanners (Epson Corporation, UK) in a temperature and humidity controlled cabinet and a consecutive series of images was produced at a periodicity of 10 minutes at 30°C for 24 hours. The lag of each growth curve was calculated as described below.

**Figure 4.**
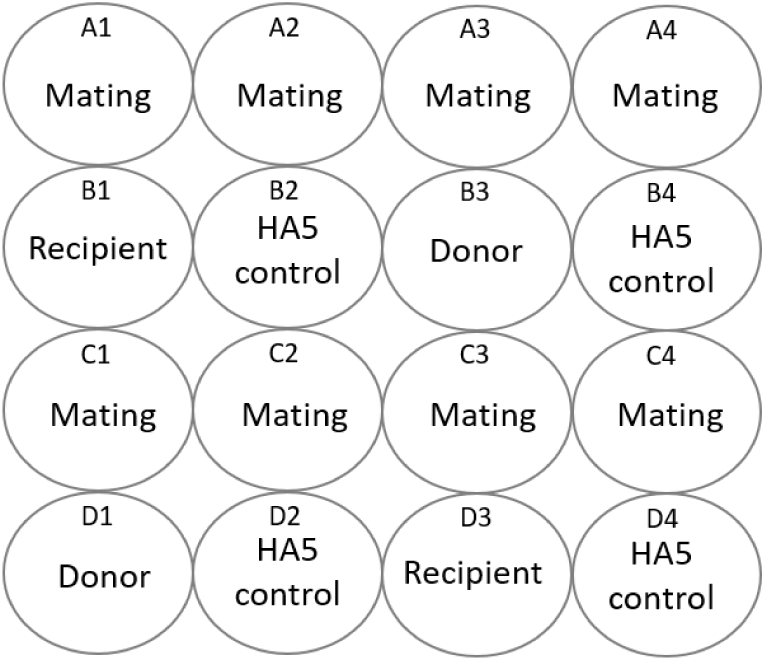
Robot pinning coordinates and pinning scheme for test of conjugation system. Pinning coordinates are identified by a letter and a number: each coordinate represents a 96-well plate and a total of 16 96-well plates can fit per plate (1536 spots in total). ‘Mating’ indicates positions were HA4 was pinned together with HA14, Negative controls are indicated by ‘Donor’ or ‘Recipient’ and ‘HA5 control’ indicates the position of the positive growth control (Tet^R^ Cam^R^). To create this pattern, three pre-culture plates were pinned together. The recipient pre-culture of HA4 was pinned from a 96 well plate into positions A1-A4, C1-C4 and B1+D3 of a LB Cam plate. Donor pre-cultures were prepared by pinning from a HA14 96-well plate to positions A1-A4, C1-C4 and B3+D1 of a LB Tet Kan plate. Positive control pre-cultures were prepared by pinning from an HA5 96-well plate to every fourth position (B2, B4, D2 and D4) of an LB Tet Cam plate. A 1536 pad was used to pin from the three pre-cultures onto a LB Tet Cam plate. The resulting format is shown and the data from this experiment is in Figure 2B.

### Construction of the donor Keio library

The Keio donor library was constructed by conjugating the F plasmid from XL1-Blue to the Keio mutants. A HDA RoToR robot (Singer LTD, UK) was utilized to construct the donor library by pinning cells from 96-well plates onto solid LB Tet Kan plates using 96-long pads (Singer LTD, UK). A 96-well plate of XL1-Blue was created by mixing an overnight culture of XL1-Blue grown in LB Tet with glycerol (20% final concentration) and pipetting 175 µl into 96-well plates. The KEIO 96 well plate was created similarly. Each Keio plate was pinned twice (positions A1 and C2, in each tetrad of positions); then the XL1-Blue 96-well plate also pinned twice (positions A1 and C3) followed by incubation at 37°C overnight. This created a mating position at A1 and negative controls for each strain (Fig 5). Only position A1 should grow as the plasmid will be transferred from XL1-Blue to the Keio mutant creating the donor strain (Kan^R^, F′Tet^R^). The following day, the transconjugant containing position A1 is pinned to position A1 of a fresh LB Tet Kan plate and allowed to grow overnight at 37°C. The purified transconjugants are pinned back to 96-well plates containing 125 µl of LB Tet Kan and incubated at 37°C for approximately 20 hours. Glycerol is added to the plates to a final concentration of 15% in a total volume of 175 µl. The Keio donor plates were frozen at -80°C.

**Figure 5.**
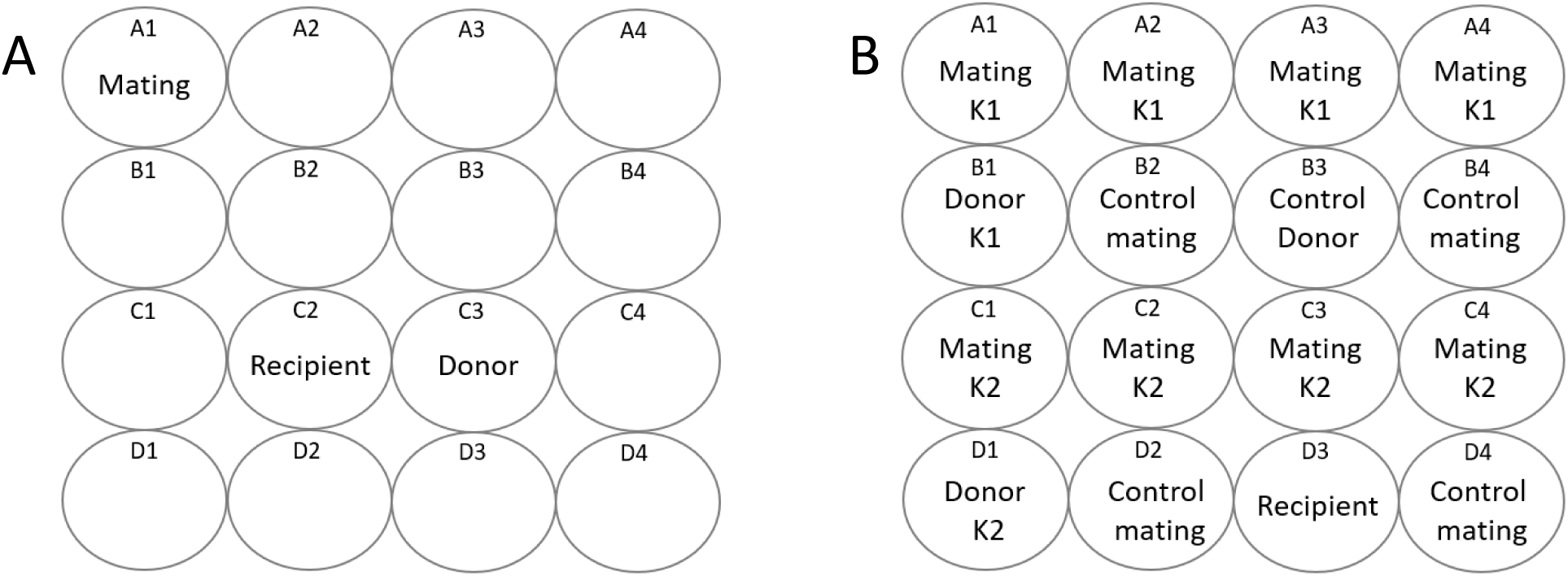
Pinning scheme for the Keio donor library construction and mating screen. **A.Donor Library Construction** Robot pinning coordinates and pinning scheme for donor library construction. The pinning scheme for donor library construction is indicated. The pinning coordinate for the mating at A1 will contain the transconjugants, while the coordinates for the recipient’s negative control (the Keio mutant) and the donor’s negative control indicated at positions C2 and C3 respectively. The remaining positions are not used. B. Donor Library Screening for Conjugation Efficiency The format of the Keio screen matings and controls are shown. Two different set of 96 Keio mutants were pinned onto one plate as indicated by K1 and K2. ‘Mating’ indicates positions where a Keio donor strain was pinned together with the recipient strain (HA4), Control matings of HA14 and HA4 are included in every fourth position and negative controls for the KEIO (Donor K1 and K2), HA14 (control donor) and HA4 (recipient) are indicated. This plate was created by first pinning strains onto two pre-culture plates and then using a 1536 pin pad to pin the final pattern onto an LB Tet Cam plate. The pre-cultures were done as follows: Donor pre-culture: positions A1-A4 and B1 are pinned with the first donor Keio plate (P1), positions C1-C4 and D1 are pinned with the second donor Keio plate (P2) and every fourth position and B3 are pinned with HA14 on an LB Tet Kan plate. On the recipient pre-culture LB Cam plate, all positions except B1, B3 and D1 are pinned with plate HA4. Example results from this experiment are in Figure 3.

### Donor Keio library screen

The KEIO donor plates were thawed at room temperature and the HDA RoToR robot (Singer LTD, UK) was used to transfer cells to an LB Kan Tet pre-culture plate in duplicate, four replicates per plate (total, *n*=8). A pre-culture plate of the recipient HA4 was made similarly. After 16 hours at 30°C the two plates were pinned together onto LB Tet Cam plates creating 4 replicate matings per plate and all appropriate negative controls were included (donors and recipients alone). A control mating of HA4xHA14 was included in every fourth position to control for any spatial variation arising from plate position. Figure 5 details the pinning scheme. After pinning the two pre-cultures together onto the selective mating plate (LB Tet Cam), the pinned plate was immediately fixed in the scanner and the experiment initiated. The secondary screen of 94 selected strains were performed similarly, with 6 replicates distributed across each of eight plates (*n*=48).

### Automated extraction of lag times

High resolution population size growth curves were obtained using Epson Perfection V800 PHOTO scanners (Epson corporation, UK) and the Scan-o-matic framework (Zackrisson, et al, 2016), version 2.0. Scanners were maintained in a single thermostatic (30°C), high humidity cabinet to minimize light influx and evaporation. Experiments were run for 24h, with automated transmissive scanning and signal calibration in 10 min intervals, as described (Zackrisson, et al, 2016). Calibrated pixel intensities were transformed into population size measures by reference to cell counts obtained by optical density measurements, using the conversion: *y* = 2.128 * 10^-2^ *x*^5^ + 1.023 *x*^4^ + 11.47 *x*^3^ + 25.62 *x*^2^. Population growth curves were smoothed to remove noise using a Lowess-like weighted polynomial function, as described (Zackrisson, 2017). Poor quality curves (0.25%) most commonly due to failed cell deposition (mis-pinning) were rejected following manual inspection. We segmented smoothed, log_2_scale growth curves to identify an initial flat phase as a sequence of at least 3 data points with the required properties -0.02<*d*<0.02, where *d* is the 1st derivative. We next segmented the remaining part of the growth curves to identify the linear phase that corresponds to the largest increase in population size. We extracted the lag time as the intercept between the initial flat and the linear phase, if the start of the linear phase occurs after the end of the initial flat phase. Details can be found in (Zackrisson, 2017; the code is available here (https://github.com/Scan-o-Matic/scanomatic/blob/1b803ab5463f027cfe106034fffc60b5b5d3a9ff/scanomatic/data_processing/phases/features.py#L417-L457).

### Confirmation of Mutant alleles by PCR

Donor strains carrying the mutant Keio alleles were analysed by standard colony PCR using the primers in Table S2 and the kanamycin cassette internal primer k1, which gives two bands if the gene has been replaced by the cassette and a single band if not, as previously described (Datsenko and Wanner, 2000). Control reactions were done on BW25113.

**Liquid mating assay:** Liquid mating assays were modified from Anthony et al 1994. Cultures of each candidate, HA14 and HA4 were grown overnight in LB with appropriate antibiotics. The following day, the antibiotics were washed off and the cells resuspended in 1 ml LB pre-warmed to 37°C. The washed donor cells were diluted 1:50 in pre-warmed LB and grown to log phase. Recipient cells were adjusted to OD_600_=3. 500 µl of each candidate or HA14 were mixed with 500 µl of HA4 and allowed to conjugate without shaking for 30 minutes at 37°C. After the incubation, conjugation was stopped by placing the cells on ice for 1 minute followed by vigorous vortexing for 1 minute then incubating on ice. Serial dilutions (10x steps) of each conjugation mixtures were prepared in 1X M9 salts (6 g/l Na_2_HPO_4_, 3 g/l KH_2_PO_4_, 1 g/l NH_4_Cl and 0.5 g/l NaCl) then 10 µl of each dilution were spotted twice on LB Tet Kan plates and LB Tet Cam and incubated at 37°C overnight to quantitate the number of donors and transconjugants, respectively. Conjugation frequency was calculated as the number of transconjugants per donor. The calculated frequency was then normalized to the mean conjugation frequency of the control matings (HA4 X HA14) and expressed as a ratio of the control.

